# Inevitability of Red Queen evolution driven by organismic complexity and simple feedback via environmental modification

**DOI:** 10.1101/2021.09.26.461893

**Authors:** Daniel S. Fisher

## Abstract

Evolution in complex high-dimensional phenotype spaces can be very different than the caricature of uphill evolutionary trajectories in a low-dimensional fitness landscape. And slight modifications of the environment can have large consequences for the future evolution. Here, the simplest approximation of evolution, an almost-always clonal population evolving by small effect mutations under deterministic “adaptive” dynamics, is studied. The complexities of organisms and their interactions with their environments are caricatured by population growth rates being smoothly varying random functions in very high dimensional phenotype spaces. In a fixed environment, there are huge numbers of fitness maxima, yet evolutionary trajectories wander around amongst saddles, gradually slowing down but still wandering widely and without committing to any maximum. But with even very small changes of the environment caused by the phenotypic changes, after an initial transient the evolution continues forever without further slowing down. In this Red Queen “phase” the apparent rate of increase of the fitness saturates (at a feedback strength-dependent rate) and the trajectories perpetually wander over large phenotypic distances. Organismic complexities, caricatured by a large number of constraints on the molecular-level phenotype, together with the simplest possible interactions of the organisms with their environment, are shown to be sufficient to cause such Red Queen dynamics. Arguments are made for the ubiquity of such behavior.

## 1 Introduction

Phenotypic evolution is often caricatured as uphill motion in a fitness landscape towards a fitness peak. But there are two essential problems with this picture. First, phenotype space is very high-dimensional and motion in complex high-dimensional landscapes is very different than in simple low-dimensional ones. And, second, “fitness” — in spite of having two ss’s — is often envisioned as a single function of the phenotype (or genotype). At the very least, the net growth rate of a clonal population is a function of the concentration of all relevant chemicals in the environment — what one might call its “fitnome”. [1] And any evolution will change the environment, even if only slightly: A snowscape, which is changed by the climber, is a better metaphor. Gradual phenotypic evolution that gradually changes the environment, and thereby also gradually changes the interactions between species, is often referred to as *adaptive dynamics*. [2, 3]

It has been shown that coevolution of several interacting species can lead to complex behavior: evolutionary cycling, [4] or evolutionary chaos, [5, 6] even in the simple framework of adaptive dynamics. Here we focus on the most basic situation, a single species, and ask: Are there properties of simple evolution in high-dimensional landscapes and snowscapes that are *general* consequences of the complexity, high-dimensionality, and even-weak feedback from environmental changes? Do evolutionary trajectories converge to a stable fixed point — albeit one of many possible? Or do they keep wandering while only gradually slowing down? Or — the most interesting possibility — can they undergo perpetual Red Queen evolution continuing to chase the changes in environment that the evolution causes? Doebeli and Ispolatov [5] —henceforth D&I — have shown that adaptive dynamics can lead to persistent evolutionary chaos. In many systems, chaotic dynamics is more prevalent in high dimensions, and D&I have shown this occurs in some models of adaptive dynamics [5, 7]. But how ubiquitous is such chaos? In particular: How much “complexity” is needed to drive Red Queen evolutionary dynamics? And, if it occurs with weak feedback, what are its characteristics?

Before developing and analyzing models, we must ask: What is the appropriate phenotypic space and its dimension? The dimension of the environmental space, *D_E_*, even in a well-mixed system with no time-dependence, is at least as large as the number of chemicals that can affect the organism. And the relevant organismic phenotype includes at least how its reproductionminus-death rate is affected by all these chemicals, and how much it produces or consumes of some of them. Thus the organismic phenotype dimension, *D_O_*, is at least as large as *D_E_*. But evolution acts, via genetic changes, most directly on the *nano-phenotype*: all the properties of the proteins and their interactions with each other and with DNA, etc. which can affect how it responds to and how it affects its environment — especially, but certainly not exclusively, via metabolic processes. This nano-phenotype dimension, *D_N_*, is very large. [8] And it is in this space of dimension *D_N_* in which all the molecular and cell-biological constraints from protein properties and prior evolution act. Thus, even if the organism interacts with its environment in a relatively simple way, with only a moderate number of environmental and organismic dimensions, *D_E_* and *D_O_*, dominating, the underlying dimension, *D_N_* is still very large. These distinctions between the nano-phenotype and organismic phenotype, and the interplay between them, are important even for simple caricature models.

### 1.1 General considerations and questions

We will denote the phenotype by a vector, **X**, in *D* ≡ *D_N_* dimensions, and the environment by a *D_E_*-dimensional vector **E**. The fitnome, Φ(**X**; **E**), is the net reproduction-minus-death rate of a clonal population with phenotype **X** in environment **E**. The environment depends on the *extrinsic* environment, **E***_ext_*, in the absence of the organisms, as well as — crucially — the population and phenotype of the occupying organisms. The simplest situation — to which we will almost entirely restrict consideration — is a clonal population at its carrying capacity in a well-mixed environment (e.g. in a chemostat). The resident population will change the environment to some 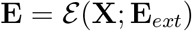: we will be interested in constant **E***_ext_* and hence drop the dependence on this. How complex functions of **X** will Φ and 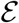 be? Any beneficial mutation in an already quite-well-adapted organism will have both positive and negative effects and these will depend on the both its evolutionary history — “genomic background” — and the environment. Thus the sign of its net effect will vary: if a mutation was unconditionally beneficial, it would already have fixed in the population. A crude caricature of the complexity is to treat Φ and 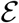 as being random functions of the phenotype **X** and characterize them by their statistics. The simplest caricature is to treat Φ at fixed **E** directly as a random “rugged” function of the nano-phenotype **X** which has very many distinct maxima. A refinement is to consider the many molecular-biological constraints — soft or hard — each as complex pseudo-random non-linear functions of the nano-phenotype, but with, at the organismic level,, Φ being a much simpler function of **X**. As we shall see, simple models based on these caricatures behave very similarly in key respects. We will thus argue that expecting substantial universality of qualitative results is not unreasonable.

We primarily consider the simplest adaptive process: a small enough population that beneficial mutations which arise sweep to fixation before any others arise, and each beneficial mutation has only small phenotypic consequences. In a fixed environment, **E**, the fitnome, Φ(**X**; **E**), is a landscape with evolution approximated by gradient ascent of the phenotype, **X**. However with the environment changed by its resident population with phenotype **X**, a mutant **Y** = **X** + *d***X** must be better than **X** in the environment 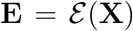. The phenotype of the almost-always-clonal population then evolves under evolutionary forces that are *not* gradients

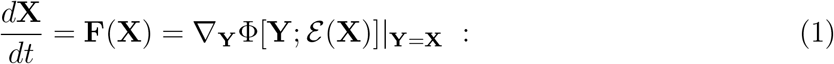

this approximation to the evolution is what-is-referred-to-as adaptive dynamics and has been extensively studied: e.g. [2, 3, 5]. Note that, for convenience, we have set the coefficient that determines the basic time scale of the evolution, to be unity: it will of course depend on many features, in particular the population size and fixation probabilities of mutations.

When there is no evolutionary feedback on the environment, an evolutionary trajectory will eventually converge exponentially to one of the maxima of Φ(**X**). But in rugged landscapes with multiple maxima, the boundaries of the domains of attraction of the maxima can be very contorted. Which maximum a trajectory converges to will be controlled by its passage near to saddle-points. Each saddle point is characterized by an index, *I*, the number of unstable directions, which will vary from 1 to *D* – 1 (with maxima having *I* = 0, and minima *I* = *D*): *I* is one property of the eigenvalues of the Hessian matrix, *∂*^2^Φ*/∂x_i_∂x_j_*, which controls the dynamics near the stationary point.

In the limit of high dimensionality, “generic” random rugged landscapes have many features that are universal [9, 10] including — crucially — exponentially many, in *D*, maxima, minima, and saddles of all indices. At any of these stationary point the eigenvalues of the Hessian will all be real with an approximately-continuous spectrum — known generically to be semicircular with its center shifted away from zero in a way that is related to its index. [9] A vast majority of the maxima are almost marginal, with their least-negative eigenvalue — the upper edge of the spectrum — very close to zero. If these marginal maxima were to dominate the late-time dynamics, generic trajectories would not converge exponentially except at very long times. Indeed, with time scaled so that the typical eigenvalues of the Hessian are of order unity, the eventual exponential convergence rate scales as an inverse power of *D*. Over a wide range of time scales before this eventual convergence, trajectories very near to an almost-marginal maximum converge towards it only as a power of time (with the power-law exponent related to the square-root edge of the spectrum of only-slightly-stable eigenvalues). But in actuality, the behavior is much more interesting: in the limit of infinite *D*, generic trajectories never “commit” to any maximum but instead wander forever amongst the saddles with Φ gradually increasing towards where the marginal maxima occur, and the index of the saddles the trajectory passes near gradually decreasing towards zero. This is known as *aging* behavior in physics contexts [11]: here the properties of evolutionary trajectories, depend on their evolutionary age. Intuition based on low-dimensional fitness landscapes is very misleading.

With the very marginal behavior of trajectories in high-dimensional fitness landscapes, it is not surprising that the behavior can be changed by even very small feedback that changes the environment slightly as the population evolves. Although the evolutionary forces, **F**(**X**), are no longer gradients, there will still be exponentially many stationary (or fixed) points [12] but with their linear stability matrices, *L_ij_* = *∂F_i_/∂x_j_*, now having some complex eigenvalues (which can cause trajectories very near a stable fixed point to spiral into it). Again, for large *D* almost all the stable fixed points will be close to marginal, and the low-index saddles will have positive eigenvalues that have small real parts.

But does small feedback qualitatively change the *global* dynamics? This is not just determined by the linearized dynamics near to saddles and stable points: the presence of very small eigenvalues means that non-linearities are very important. We are particularly interested in the behavior at long times. Does the evolution gradually slow down, as in a fixed landscape? Or does the rate of evolution converge to a non-zero feedback-magnitude-dependent rate at long times? How, and for how long, does the evolution depend on past history? Quantitatively, the rate of evolution can be characterized by the *fitness flux* [13]: the rate of change of the fitness measured with respect to the *current* environment,

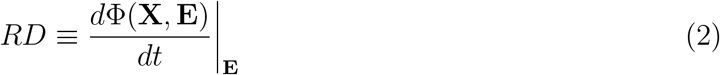

which for adaptive dynamics, is *DR* = |*d***X**/*dt*|^2^; we have included a factor of *D* to make *R* of order unity. A natural definition of Red Queen behavior is fitness flux not tending to zero at long times. Other quantities of interest include the degree of persistence of trajectories at short times, their longer-term sensitivity to perturbations, and the dependences of the characteristic time scales on the strength of the environmental feedback.

## 2 Models and Approach

To make analysis tractable, we consider random-landscapes with simple statistical properties. To keep the phenotypes from wandering arbitrarily far, we confine **X** = {*x_i_*}_i=1,2,…*D*_ to lie on a *D*-dimensional (strictly speaking *D* – 1 dimensional) hypersphere with 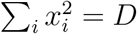 so that the {*x_i_*} are of order unity. (It is known that other ways of confining trajectories, e.g. [9, 14] yield very similar behavior.)

### 2.1 Gaussian random fitnomes

We consider ensembles of gaussian random fitnomes: specifically taking the forces **F**(**X**) to be gaussian with mean zero with covariances that depend only on the angle between two points: **X** and 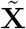: i.e. the ensemble is statistically rotationally invariant in **X**. In the absence of environmental feedback (when 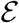 is independent of **X**) we take the landscape Φ(**X**) to have mean zero and covariance:

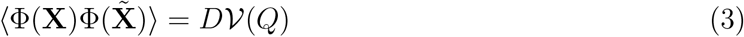

with

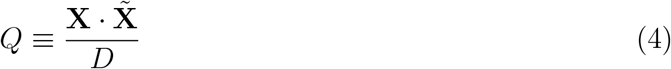

so 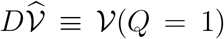 corresponds to the variance and we have introduced the notation that the local quantities which are parametrized by *Q* =1 — i.e. 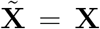 and later derivatives and curvatures — are indicated by hats. The covariance can formally be expanded in integer powers of *Q* and, for *D* → ∞, the condition for being a legitimate covariance is that all the coefficients of this expansion be non-negative. [15] The evolution is simply gradient ascent on this landscape, with an **X**-dependent, and therefore implicitly time-dependent, Lagrange multiplier, *μ*(*t*) = **X**(*t*) · ∇Φ/*D*, needed to remain on the hypersphere:

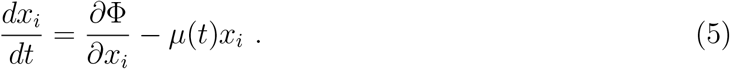

The covariances of the gradient forces are:

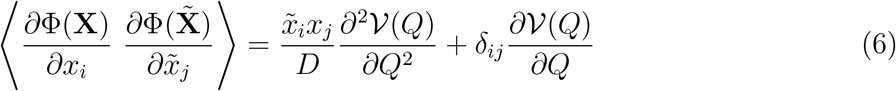

with the scalings with *D* making the force that acts on a single component, *x_i_*, be of order unity.

In the presence of feedback, there will be additional forces that are generally not gradients of a potential. The simplest form to assume for the covariances of the additional feedback forces,, is just proportional to *δ_ij_* like the second term in Eq.(7). (The more general form that is still statistically rotationally invariant will also have a 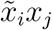 part but this can be included in a redefined 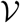.) We denote the magnitude of the feedback forces as 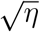 so that the additional covariance is proportional to *η*, and write the covariances of the total evolutionary forces as

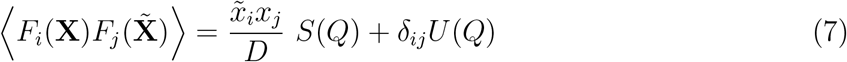

with again 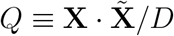,

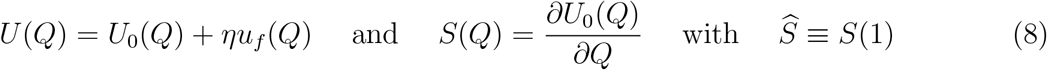

setting the basic time scale: 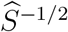. The covariance of the feedback part, *u_f_* (*Q*), should also be a sum of powers of *Q* with non-negative coefficients.

#### 2.1.1 Toy *p – H* model

A particularly simple family of fitness landscape models is

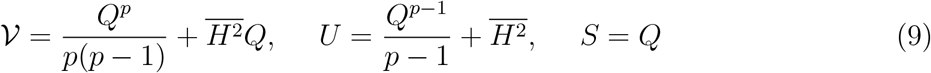

for *p* ≥ 2 an integer, and normalization chosen to make 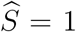 for convenience. We will refer to this class of models for *p* ≥ 3 as “*p* – *H* models” and for *H* = 0 as “pure-*p* models”. (In the context of spin-glasses for which they were developed, these are referred to as “p-spin spherical spin-glasses in a field”.) [16, 17] For *p* =3, which we will use for numerics, the fitness is 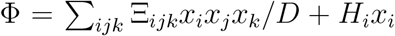 with the 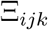 iid gaussian, and 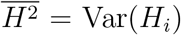. The “field” — or *bias* — gives a preferred phenotype direction: if 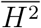 is large enough, there will be only one fitness maximum.

Feedback can be added simply by, for example, multiplying the *Q*^*p*–1^ part of *U* by (1 + *η*). Other forms of feedback will behave similarly as long as *u_f_* (*Q*) depends on *Q*.

In the special case, *p* = 2, a matrix — with feedback a non-symmetric matrix —parametrizes the interactions. For 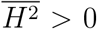, this model behaves quite simply. Direct analysis is enabled by the linearity of the dynamics except for the constraint reflected in the time dependence of *μ*(*t*). We quote some results for this matrix model in Secs. 3.0.1 and 4.1.1.

### 2.2 Modeling constraints

An alternative class of models — much better motivated biologically — is to separate the nanophenotype and the organismic phenotype and approximate the fitnome as the sum of two parts: a complex function of the high-dimensional nano-phenotype, Φ*_cnstr_* (**X**), which does not depend on the environment, and a simpler lower-dimensional function of the organismic phenotype and the environment, Φ*_envt_*(**X**, **E**). The feedback from modification of the environment only depends on the organismic level phenotype but this of course depends on the nano-phenotype, even if in only a simple way. The genetically determined nano-phenotype is very high-dimensional and the organism’s fitness will strongly depend — in a largely environment-independent way — on all the genetically determined quantities. Most simply, this could be modeled as a large number of molecular and cell-biological constraints with the basal fitness, Φ*_cnstr_*, being constant over the portion of the nano-phenotype space over which all the constraints are satisfied.

As a caricature of the molecular level interactions, we consider them as imposing a large set of *K* constraints. These constraints we write in the form Γ*_α_*(**X**) = 0 for each of *α* =1, 2, … *K* and define *κ* = *K/D*. More generally, the constraints could be soft and we caricature the associated fitness cost whose gradient contributes to the evolutionary forces by a parameter Ω: Ω → ∞ gives rigid constraints. For rigid constraints, one must certainly have *K* < *D*; however, the condition for all the constraints to be simultaneously satisfiable can be stronger than this, Sec.5.1.1. In the under-constrained regime, there is no cost to phenotypic changes on a neutral *D* – *K* dimensional sub-manifold of the phenospace. With Ω finite the system could be over-constrained, there then always being an associated cost from not being able to satisfy all the constraints: we will not investigate that regime in detail.

We model the constraint functions {Γ_*α*_} by gaussian random functions with mean zero and covariance

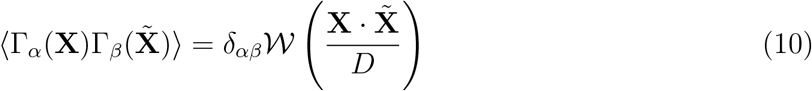

and the fitness cost of violating the constraints by

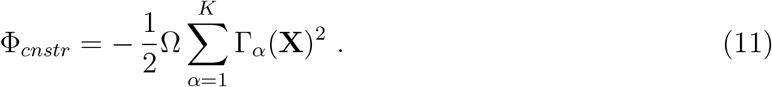

It will turn out that minimally non-linear constraints — simply pairwise “interactions” — are sufficient to demonstrate rich behavior. These can be caricatured by

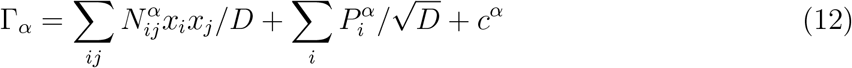

so that

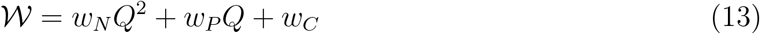

with non-negative coefficients. If the *c^α^* are large — highly *skewed* constraints — then it can become impossible to satisfy all the constraints even with *κ* < 1: indeed, with *w_N_* = 0 and *w_C_* > 0, this will always occur. The analytical results we derive are for general 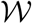 but for numerical calculations we primarily take the form in Eq. (12) simplifying further to *w_P_* = *w_C_* = 0.

### 2.3 Simplest coupling to environment and feedback

At the level of the *organismic* phenotype, interaction with a simple environment can be parametrized by a modest-dimensional vector, **E** = {*E_ρ_*}, with *ρ* = 1, 2, … *D_E_*. The simplest characterization of the effects of a population on the environment and the effects of the environment on the population is via a pair of *D_E_* dimensional vector functions of **X** that characterize, respectively, the effects of the population of phenotype **X** on the environment and the changes in its population growth rate caused by environmental changes. With respect to some reference environment, **E**^0^, set by the combination of external conditions and a saturated population of some reference phenotype, we assume only small changes, **E** = **E**^0^ + Δ**E** and model the population growth rate as simply 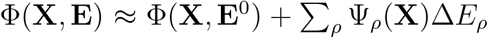. To avoid any complexity in the dependence on the nanophenotype, we assume a linear dependence on changes in the phenotype (loosely as if expanding around the reference phenotype): 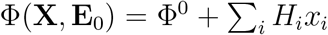 and 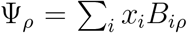, with *H_i_*(**E***_ext_*) depending on the extrinsic, externally controlled aspects of the environment, and we have absorbed the constant parts which do not affect the dynamics. Similarly, we assume that the effects on the environment depend linearly on the nano-phenotype: 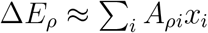. Putting these together we have for the organismic level fitness of phenotype **Y** in an environment set by a population of phenotype, **X**, simply:

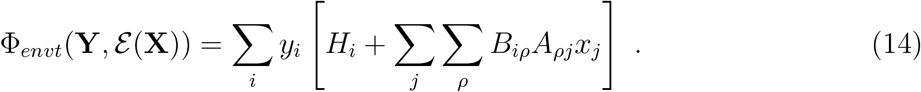

The evolutionary forces from the interactions between the organism and environment are then simply of the form:

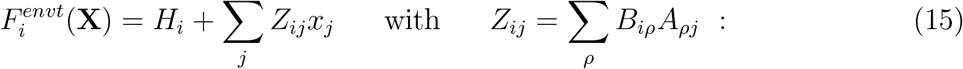

**Z** is thus a rank-*D_E_* matrix. with *D_E_* ≪ *D* ≡ *D_N_*.

The simplest situation is if **A** and **B** have *independently* gaussian random elements. Even then, **Z** is not gaussian. However the deviations of its elements from gaussian are small for even modestly large *D_E_* — i.e. a relatively simple environment — and their covariance can, for the properties needed in the analysis, be well-approximated by treating the elements *Z_ij_* as if they were independently gaussian random with variance *η/D* — this variance thereby defining *η* and the covariance function of the feedback forces simply *ηu_f_* = *ηQ*. In the Discussion, we consider effects of correlations between **A** and **B** as would occur if, for example, the changes in the environment were due to consumptions of resources [18])

The main goal of the simple caricatures of the interactions with the environment is avoiding complexities or multiple maxima in the dependences on both the nano-phenotype and the environment, not in the linearization *per se*. The crucial feature that drives the interesting evolutionary dynamics is that the phenotypic dimension, *D* = *D_N_*, is very large and the nonlinear constraints make the fitness as a function of the nanophenotype very complex — albeit in a way that neither substantially effects, nor is substantially affected by, the environment.

### 2.4 Dynamical mean-field theory

With gaussian distributions of the evolutionary forces or gaussian constraints with simple covariances, in the limit of high dimensions analytic progress can be made via use of dynamical mean-field theory (DMFT) developed for studying spin-glasses and other random physical systems. [16, 11, 19] The crucial enabling property is that the effects of each of the many other component of **X** on one component of interest, *x_i_*, act like a gaussian random force on it — with correlations in time reflecting the dynamics of the other components. But, in addition, a change in *x_i_* causes small changes of each of the other *x_j_* which result in changes in the forces back on *x_i_* — a consequence of the correlations between *∂F_i_/∂x_j_* and *∂F_j_/∂x_i_*. This effect is proportional to the averaged response to a small perturbation of the forces on the other *x_j_*. The effective dynamical equation for *x_i_* is then:

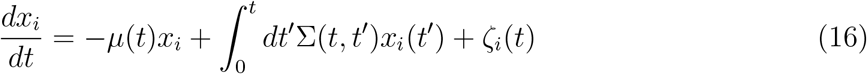

with self-consistency — below — forcing Σ to be a specific functional of the correlation function

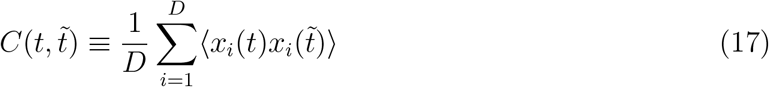

and the (average) response function

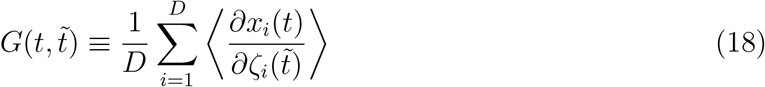

to a small addition force applied at time 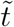 (so that *G* = 0 for 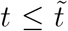). The averages are taken over the gaussian “noises” ζ*_i_*(*t*) and the normalization of **X** fixes *C*(*t, t*) = 1 for all *t*.

The dynamical equation (16) can be multiplied by 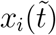 and then averaged over the noise to yield:

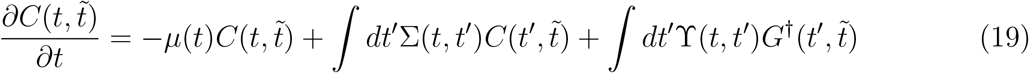

where the dagger is like a matrix transpose: 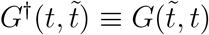. Since as a function of *x_i_*(*t*) the dynamical equation is linear, the response function can be found directly by taking a derivative with respect to the noise yielding

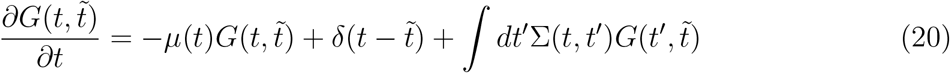

As the noises arise from the dynamics of the other components, and the feedback on the component of interest involves correlations in the forces, two self-consistency condition must be obeyed. For the simple model in which the evolutionary and feedback forces are gaussian random, Sec. 2.1, we have

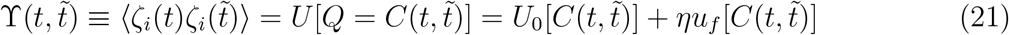

and

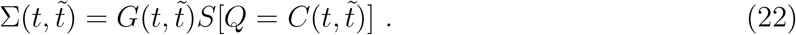

Importantly, note that, since *S* = *dU*_0_/*dQ*, Σ does not have a contribution from the feedback part (or more generally has only a smaller contribution) because it does not arise from gradient forces.

We will sometimes, for shorthand, denote time integrals such as occur in Eqs. (19,20) as if they were convolutions (which they are if the functions only depend on time-differences): e.g. writing ϒ ⊗ *G*^†^ (for discrete time, this is just matrix multiplication). In the model with constraints, the DMFT equations have exactly the same form, but Σ and ϒ become complex functionals of *C* and *G*.

## 3 Random landscapes with no feedback

Much is known about dynamics in static high-dimensional random landscapes and we can make use of these results. (In most analyses, “thermal noise” is added to the dynamics: we are primarily interested in the behavior of deterministic dynamics — i.e. zero temperature — but the behavior is similar to that at low temperatures. [16])

There are two main classes of high-dimensional random landscapes with covariances of the forms we have assumed. The simplest is characterized by a unique maximum: this occurs trivially if Φ = **H** · **X**, corresponding to 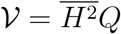. But small additional non-linear contributions to 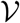 do not change the behavior — for large 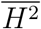 the toy *p* – *H* model [Eq.(9)] exhibits only a single maximum. As when there is only one or a modest number of stable fixed points, the trajectories will converge to one of these, we call this *convergent* evolutionary dynamics.

The other generic class is landscapes with exponentially many maxima. Although there are several sub-classes (known to cogniscenti as ‘1-step-RSB” and “full-RSB”) [20] but for most of our purposes they behave similarly. [10, 19]

### 3.0.1 Matrix model

In addition to the two main classes, there are special non-generic cases with a modest number of maxima, most simply when there is a symmetry. A very special case is with two-fold symmetry and particularly simple interactions: 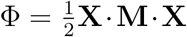 with **M** a random matrix (the *p* – *H* model with *p* =2 and 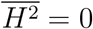). There are two equivalent maxima corresponding to **X** aligned parallel or anti-parallel to the eigenvector of the largest eigenvalue of **M**. And there are many saddle points (corresponding to other eigendirections of **M**) which make the dynamics from random initial conditions interesting — although, except for the Lagrange multiplier, the dynamics is linear. The abundance of saddles still gives rise to slow aging dynamics as discussed in Sec.3.2. [21] But any small perturbation destroys the special structure: in particular, with a small **H** · **X** term added to Φ, there will immediately be only one maximum and (unless the components of **H** are smaller than an inverse power of *D*) the saddle points will also be destroyed. [21]

### 3.1 Convergent dynamics

When there is a unique maximum, the behavior is simple: trajectories converge exponentially and the response to a perturbation applied after a long time has effects that decay exponentially with time. At long times, the correlation function, 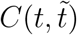, approaches unity, for all time differences

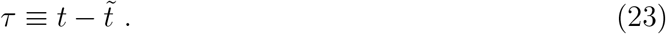

The response function for *τ* ≪ *t* can then be found directly by Laplace transform yielding:

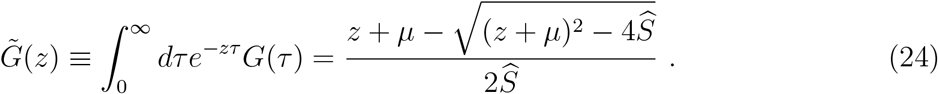

This reflects, in the imaginary part of 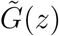 as *z* → λ, a semi-circular density of states centered at λ = –*μ* with edges at 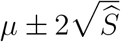. When there is a single maximum the eigenvalue spectrum of the Hessian has a gap with 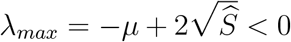 which controls the exponential approach of trajectories to the maximum. With *G*(*τ*) decaying rapidly and *C* ≈ 1, the integrals in the 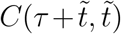 equation (19) can be immediately done to yield an equation relating the *susceptibility* 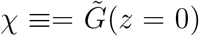 — which characterizes the net response to a perturbation — to *μ*, 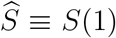, and 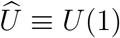: simply 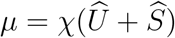. Together with Eq.(24), this yields

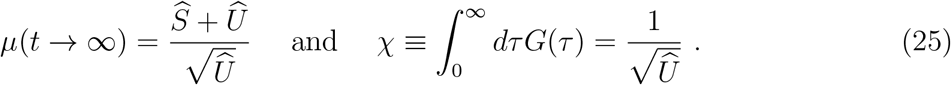

To be consistent with being at a maximum, i.e. λ*_max_* < 0, one thus must have that 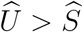: this is the condition to be in the *convergent phase* with a unique maximum and the concomitant exponential convergence of trajectories towards it. For the *p* – *H* model, the critical squared bias is 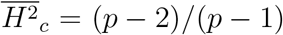. (For the matrix model, *p* =2, this recovers the result that the 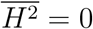 model is special, including being exactly critical.)

In the convergent phase, memory of the past decays away so at long times the projection of **X** on the direction of **H** can be found simply: **X**(*t* → ∞ · **H**〉/|**H**| = χ|**H**|. However **X**_∞_ also has components perpendicular to **H** due to the effects of the random interactions.

An understanding of the condition for the self-consistency of the convergent phase can be obtained, more generally, by adding a small noise at frequency *ω* with power spectrum 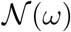. Near a maximum with a gap in its spectrum this should only induce small fluctuations in **X**. Adding this external noise into the DMFT equation for *C* and expanding around *C* = 1, shows that this external noise induces, via the interactions between the components, an effective noise on each *x_i_* with correlation *δ*ϒ(*ω*). The linear response to the additional noise will result in 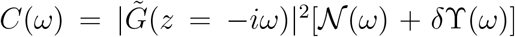. But this change in *C* corresponds to a change in ϒ, proportional at low frequencies to *dU/dQ*|_*Q*=1_, which for gradient forces is exactly 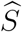. Self-consistency thus mandates that in the limit of low frequencies, 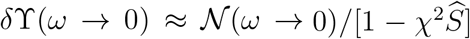. But from Eq.(25), 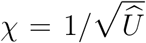 which implies that the non-linear response that determines *δ*ϒ diverges as 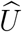 decreases to 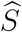, indicating failure of the assumption of a stable maximum.

We will later see that that for non-gradient dynamics the two conditions derived above are *not* the same: the noise amplification diverges while the averaged response function, *G*, is still non-singular.

### 3.2 Many maxima, no feedback: aging dynamics

When there are exponentially many maxima, the fitness initially increases rapidly, but then slows down gradually approaching an asymptotic value which is the fitness at which almost-all the borderline maxima occur. The fitness approaches this level, Φ_∞_, as an inverse power of time [16] (although more complicated behavior can occur for some 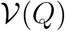. [19]) Because of this slowing down, after waiting a long time, 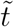, the correlation function for 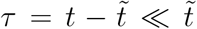 only deviates slightly from its equal-time value of unity: for 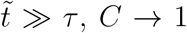, *C* → 1 indicating no motion. But the linear-response function approaches a non-trivial limit: 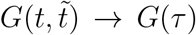 for 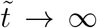 at any fixed *τ*. Its Laplace transform is still given by Eq.(24), but now we expect, since the maxima and low-index saddles that the trajectories approach near to become closer-and-closer to marginal, that

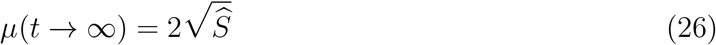

so that 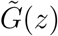 has a 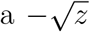 singularity for small *z*. This corresponds to power-law behavior for 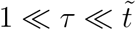:

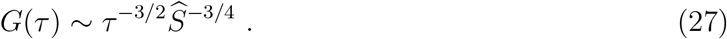

But only some of the behavior is captured by this part of the response function which probes the vicinity of the current **X** on time scales faster than **X**(*t*) moves significantly.

For *t* and 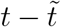 both large but of the same order, the trajectories show much more complex behavior reflected in the aging of the correlation and response functions. In the toy pure-*p* model, *C* becomes a function of 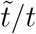 so that after any (even very long) time, 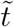, if the trajectories are followed further, up to, e.g. 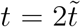, then from 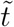 to *t* the trajectories will have wandered a distance that is a substantial fraction of the diameter of the phenotype space. Associated with this wandering, there is an *additional* slow part of the response function of the form 1/*t* times a function of 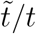. [16]

The slowing down of the trajectories results in the fitness flux gradually decreasing:

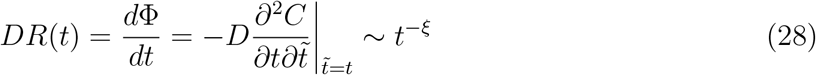

corresponding to the fitness approaching its infinite time limit, Φ_∞_, as 1/*t*^ξ–1^. In contrast to the shortish time response, the exponent ξ depends on the whole evolutionary history — via *G* and *C* for long times — and hence the landscape statistics 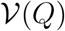. One reflection of this, is that even after a long time, 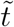, the integrated response of a perturbation applied at 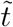,, 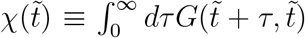 has comparable contributions from the fast part Eq.(24) with 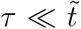 and the slow part with *τ* of order 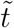: the former is controlled by the curvatures near to 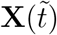 and the latter by the long term effects of the perturbation on the future trajectory. This effect is large because the trajectories continue to wander over much the phenotype space — in spite of continually slowing down.

For the aging phenomena in a fixed rugged landscape, there are several classes of behavior, depending on the statistics of the landscape, 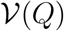, [19] but the subtle differences between them will not concern us here.

## 4 Random fitnomes with weak feedback

We now add environmental feedback with typical magnitude 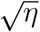. The DMFT equations have the same form except that now 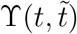 has a small extra part, 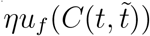, without the corresponding part in Σ from *S*.

### 4.1 Destruction of a stable maximum by feedback

Around a general stationary point with feedback, the linearized dynamical matrix, *L_ij_* = *∂F_i_/∂x_j_*, will not be symmetric and thus have some complex eigenvalues. Analyzing the stability via DMFT is straightforward, as carried out in Sec. 3.1.

#### 4.1.1 Matrix model with feedback

To understand the behavior, it is instructive to first consider the simple matrix model, Sec. 3.0.1, for which one can analyze the stability matrix directly; the dynamics are linear, 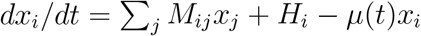, except for *μ*(*t*), but *μ* will be constant at a fixed point. For 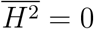, the matrix model is exactly marginal with aging, as in the absence of feedback.[21]. But for 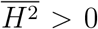 the matrix model has a unique stable fixed point (the only other stationary point is fully unstable) and its stability matrix is *L_ij_* = *M_ij_* – *μδ_ij_* (projected onto the hypersphere) whose largest eigenvalue must be non-negative: i.e. *μ* must be larger than the real part of the largest eigenvalue of **M**. With **M** random with mean zero and covariances 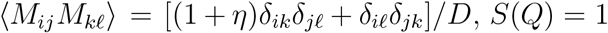 and 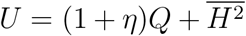. The distribution of eigenvalues of **M** is uniform inside an ellipse in the complex plane extending out to 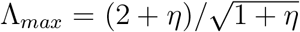 along the real-axis and up to 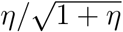 along the imaginary axis.

The analysis of Sec.3.1 gives 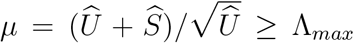 so that *μ* decreases to Λ*_max_* as 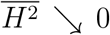. Thus for positive 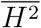 the fixed point is fully stable, while at 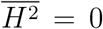, *μ* = Λ*_max_* correctly indicating the marginality. But the Laplace transform of the response function, 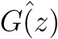 — here just the averaged Greens function of the random matrix *L_ij_* = *M_ij_–μδ_ij_* with eigenvalues λ = Λ – *μ* — does *not* appear to be marginal: it has a branch cut at 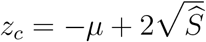 which is negative even at 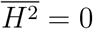. This is an artifact of the averaging over randomness: in the infinite *D* limit the naively-computed averaged Greens function for random non-symmetric matrices does *not* exhibit the singularity at the edge of the density of states. However the correct stability condition is found by analyzing the mean-square response to a small added noise, as in Sec.3.1: the stability condition is that 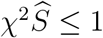 which corresponds exactly to *μ_min–stable_* ≥ Λ*_max_*.

#### 4.1.2 Feedback-driven transition out of convergent phase

For more general models with feedback we can again analyze the nonlinear effects of noise in the convergent phase in which there is a unique maximum. This gives the condition for the dynamics to converge to the stable fixed point with *C*(*τ*) = 1, and 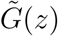 given by Eq.(24). The condition is simply 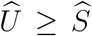 which, by analogy with the above analysis of the matrix model, corresponds to the condition that the real part of the largest eigenvalue of the stability matrix about this fixed point is Re(λ*_max_*) ≤ 0. As *η* is increased 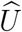 will increase until a critical value, *η_c_*, at which the fixed point goes unstable and the convergent phase breaks down.

For a concrete example, the *p* – *H* model with *p* = 3 and feedback of similar form has 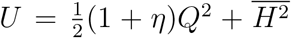. The critical 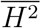 at *η* = 0 is 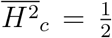 while for 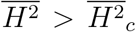 there is a non-zero critical *η*, 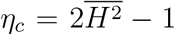. For *η* > *η_c_*, the Red Queen phase — analyzed below — occurs, while for *η* < *η_c_*, there is a unique stable fixed point and the feedback does not change the behavior qualitatively (beyond possibly causing trajectories to spiral into the fixed point).

### 4.2 Red Queen phase: complex landscape with weak feedback

We now come to the key question: In the phase in which there are exponentially many maxima of the landscape, what are the effects of *weak* feedback? There will still be exponentially many stable fixed points — those with a gap in the spectrum of their Hessian will remain stable with weak feedback — but their basins of attraction will, collectively, occupy only an exponentially small fraction of the space. The far more abundant fixed points that were marginal maxima without feedback, are highly sensitive to the small changes caused by the feedback: a linear analysis thus has very limited regime of applicability.

We make the Ansatz that for arbitrarily small *η* (but not as small as an inverse power of *D*) and almost-all initial conditions (in the large *D* limit), the dynamics converges at long-times to a strange attractor: a deterministically chaotic state. This Red-Queen phase is a *statistical steady state* of the high-dimensional dynamics with *C* and *G* functions of only time differences 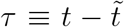. We analyze the behavior in the small *η* limit, show that the Ansatz is correct for the simplest models, and then argue that it is correct for the constraint models and far more generally.

We will show that for weak feedback there are three important time scales characterizing the steady state dynamics. The fastest time scale is set by the curvatures of the potential and is of order 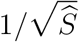 independent of *η* (we will refer to this scale as order unity). But after the initial transient the trajectories will only move slowly and, on this fast time scale, without much change of direction. Trajectories will persist in roughly the same direction for a characteristic persistence time, *τ_persist_*, after which non-linearities become important and trajectories change direction driven by proximity to saddles. For small *η*, we will show that *τ_persist_* ~ *η*^−*γ*^ and during a persistence time, the trajectories typically move a distance of order *η^ϵ^* with the exponents *γ* and *ϵ* to be determined. On time scales much larger than *τ_persist_*, the trajectories wander around seemingly randomly — from the deterministic chaos — until after a much longer *wandering time, T_W_* ~ *η*^−ζ^, with *ζ* > *γ*, trajectories have wandered over much of the hypersphere.

For *τ* ≪ *T_W_*, the correlations *C*(*τ*) will be close to unity and will not show interesting features on the fast time scale. But the response, *G*(*τ*), to a perturbation will have a fast part driven by the curvature of the landscape which is similar to that near marginally stable maxima in the absence of feedback. On longer times scales, as the trajectory moves, the response to an earlier perturbation will change because of the non-linearities. We thus make the Ansatz that the response can roughly be separated into a fast and a slow part: *G* ≈ *G_F_* + *G_S_*. How these link together on the intermediate time scale, *τ_persist_*, is subtle, as we shall see.

#### 4.2.1 Fast response and persistence of trajectories

The fast part of the response, *G_F_*(*τ*), is approximately given by the inverse Laplace transform of Eq.(24) in the close-to marginal limit, 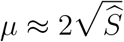. At short times this response is determined by the semi-circular density of eigenvalues of the Hessian, and over a range of time scales much larger than 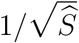, the square-root singularity at edge of the spectrum makes the response function decay as *G* ~ 1/*τ*^3/2^, Eq.(27). But this power-law form will be cutoff — by the effects of non-linearities — on the persistence time scale.

The short-time persistence of trajectories over times 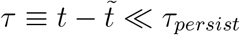, is reflected in the deviation of the correlation function from unity, which we denote Δ:

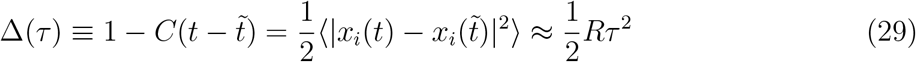

as the fitness flux is equal to the mean-square speed in pheno-space: *DR* = 〈|*d***X**/*dt*|^2^〉. The fitness flux must depend on the deviation from the gradient flow parametrized by *η*. In steadystate it is self-consistently determined by the exact expression

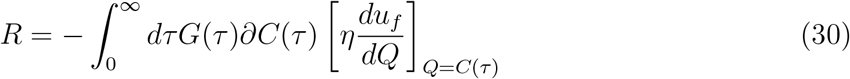

with *∂C* ≡ *dC*//*dτ* ≤ 0 and the term in brackets is 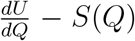 which vanishes, as it must, without feedback. (Of course, if *u_f_*(*Q*) were constant, there would be no actual feedback.) The slow increase of –*∂C* from Eq.(29), means that the short-time part of the integral for *R* is small in spite of the large response, *G_F_*, in this regime. Thus the integral is *not* dominated by short times: indeed, it can have a major contribution from the very long times of order *T_W_*, to which we now turn.

#### 4.2.2 Slow wandering dynamics

The slow parts of *G_S_* and *C* vary on the slow wandering time scale, *T_W_*, which is much longer than the persistence time. On this long time scale, the time-derivatives of *C* and *G* in Eqs.(19,20) are negligible compared to the integral terms. And the integrals over the fast part of *G_F_* act like a delta-function with amplitude 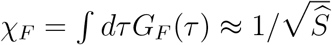 which is the net fast response to a kick.

Examination of the structure of the coupled *G_S_* and *C* equations, suggests a solution of the slow dynamical equations of the form

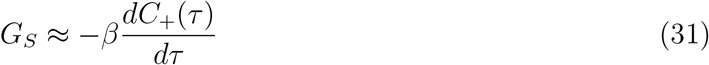

with the “+” denoting restriction to *τ* > 0 and the coefficient, *β*, yet to be determined.

Surprisingly, the coefficient *β* which relates the slow parts of the response and correlations in a way that is like the inverse temperature in a thermal system (the fluctuation-dissipation theorem) turns out to be independent of the magnitude of the feedback for small *η*: one might have expected that the effective “temperature” would be very small in the limit of small feedback. But with *β* of order unity, the form for *G_S_* implies that the total integrated response, χ, has a contribution from the slow part that is comparable in magnitude to that from the fast part.

The slow integral equation for *C* is, for positive *τ*,

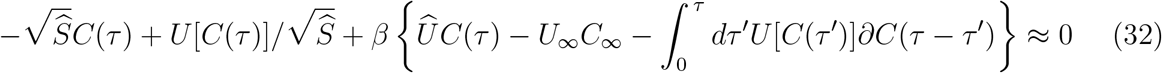

where we have used that for *η* = 0, *S*(*Q*) = *dU*_0_(*Q*)/*dQ*, to cancel the integral over longer times that appears in Eq.(19). In the presence of an **H** · **X** term in Φ, each component will have a non-zero steady-state average, 〈*x_i_*〉, so that

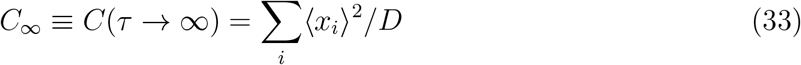

is non-zero, as is *U*_∞_ ≡ *U*(*C*_∞_).

The correctness of the Ansatz for *G_S_* can be seen by taking a derivative of the slow *C*(*τ*) equation, (32), which yields the slow equation for the symmetrized combination 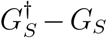. But the wandering time scale, *T_W_*, is at this point arbitrary. Indeed, the slow dynamical equation is time-scale invariant as well as being independent of *η* for small *η*. [Such time-scale invariance was first found for the aging behavior of the *p* – *H* model with thermal noise but no feedback. And the slow parts of the (there-non-equilibrium) response function were found to be similarly approximately proportional to the derivative of the correlation function. [16, 11]]

An interesting feature of the Red Queen phase in these models is that, while the time scales depend on the magnitude of the feedback, the *shapes* of the slow parts of the response and correlations become independent of *η* for small feedback. And the above results illustrate an important aspect of the eco-evo dynamics: after a long time even the near-term future depends on the long time evolutionary history, here characterized by *C* and *G_S_* with aspects parametrized by *β*. The neighborhood of a trajectory in the Red Queen phase is *not* statistically similar to a typical phenotype with the same fitness. (Because the feedback is only weak, the “fitness” is still approximately well-defined for small *η*.) Such dependence of evolutionary trajectories and local landscape on the long-time past evolutionary history has also been studied in genotypic landscape models. [22]

#### 4.2.3 Slow dynamics of pure-*p* model

In general, the slow integral equation, Eq.(32), cannot be solved in closed form. But for the pure-*p* model, 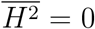 so that *C*_∞_ = *U*_∞_ = 0 and the power-law forms of *U*_0_ and *S* are simple enough to enable guessing a simple solution for the long time scale dynamics:

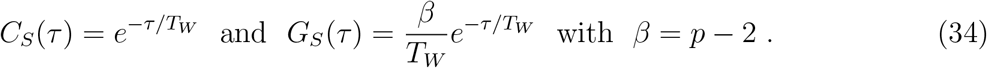

Note that, as anticipated, for *p* = 2 there is no Red Queen phase.

#### 4.2.4 Effects of bias and boundary of Red Queen phase

If a bias towards particular phenotypes, 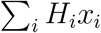, is added, then at long times the correlation function will not decay to zero as 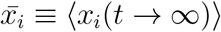 will certainly have a component proportional to *H_i_*. But every other component, 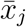, induces, via the interactions between components, a random extra persistent force on *x_i_*. Thus 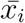 will not be simple. However in the Red Queen phase the effect of the initial conditions decays exponentially and the *time-averaged* effect of *H_i_* on *x_i_* can be found using the linear response, yielding 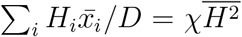. The correlation function for large *τ* is given by the 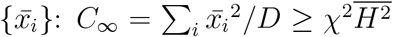, the inequality because the contributions of the mean square of the induced random components of 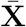 perpendicular to **H**. In the Red Queen phase, the trajectories fluctuate a lot decreasing the 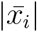 and hence making *C*_∞_ < 1 even for *η* → 0. In general, we expect the chaos driven by the feedback to make *C*_∞_ decrease further with increasing *η*.

For general models — including the *p* – *H* model with 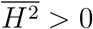 — the slow-dynamic equation (32) cannot be solved. But the quantity *β* together with *C*_∞_(*η* → 0) can be obtained without knowing *T_W_*, from the *τ* = 0 and *τ* → ∞ limits of the slow equation Eq.(32). These yield an implicit equation for *C*_∞_ in the limit of small *η*:

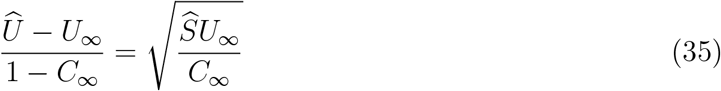

with the ratio of *G* to –∂*C* in the slow regime given by

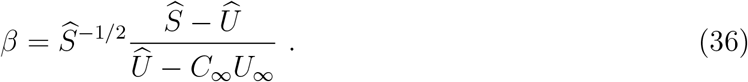

We can now see how the Red Queen phase disappears as 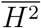 increases — or more generally as 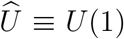 increases — for small *η*. The range over which the trajectories wander, roughly 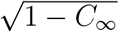, decreases until the critical point where the Red Queen dynamics ceases. Since for *η* → 0, *S* = *dU*_0_/*dQ* → *dU*/*dQ*, we see that *C*_∞_ ↗ 1 for 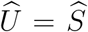: the same condition for the transition that we found coming from the convergent phase in Sec. 3.1. As this critical point is approached, 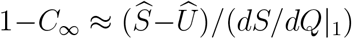 vanishes linearly but the scale 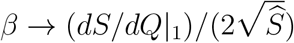 is non-singular.

At fixed small *η*, there is of course only a crossover as 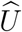 increases through 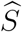 since the Red Queen phase persists slightly past 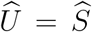 for non-zero *η*, Sec.4.1, disappearing at a critical value of 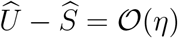.

#### 4.2.5 Matching fast to slow regimes on persistence time-scales

For the pure-*p* model, we see that in the limit *τ* ≪ *T_W_* the slow-time-scale approximation to the correlation function, Eq.(34), does not match onto the dynamics on short time scales, Eq.(29): the former has a cusp while the latter is smooth. Thus the fast and slow parts of the dynamics must splice together over an intermediate range of time scales, the natural Ansatz being *τ_persist_*, with 1 ≪ *τ_persist_* ≪ *T_W_*. To understand this matching, we start by considering the beginning of the slow-time regime for general models. For times *τ* ≪ *T_W_*, *C* will be close to unity. Thus we can expand the dynamical equation in powers of Δ ≡ 1 – *C*. The general slow equation becomes, to lowest non-trivial order in Δ,

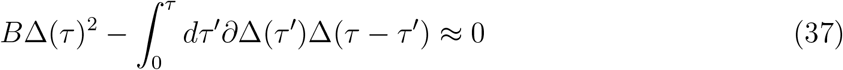

where we have normalized time scales by choosing 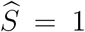 and the coefficient is then *B* ≡ *dS*/*dQ*|_1_/(2*β*). This equation is immediately seen to have a family of solutions — corresponding to the time-scale invariance of the slow dynamics — of the form

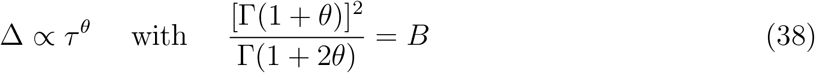

determining *θ* as a function of *B*. Since *β* depended on the whole range of the slow dynamics, Eq.(36), so does the exponent *θ* which is ≤ 1.

One might have expected that on time scales much longer than the persistence time (but before the curvature of the sphere is felt) that the randomness would make the trajectories roughly diffusive. But this is wrong: the behavior is generally *sub-diffusive*. The exceptions are the pure-*p* models: for these, *B* = 1/2 so that *θ* = 1, corresponding to Δ = 1 – exp(–*τ*/*T_W_*) ≈ *τ*/*T_W_*. As 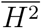 is added to *U*, and more generally if 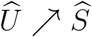, *B* increases, approaching unity at the transition as derivable from Eqs.(35,36). Concomitantly, *θ* decreases towards zero at the transition out of the Red Queen phase.

Missing from the slow-time analysis is the — still unknown — relationship between the wandering time scale and *η*, conjectured to be of the form, *T_W_* ~ *η*^−ζ^. Indeed, the scaling of the persistence time, conjectured to be *τ_persist_* ~ *η*^−*γ*^, and the persistent distance, *δx_i_* ~ *η^ϵ^* are also unknown: *γ*, *ζ*, and *ϵ* must be determined by matching the regimes.

On the persistence time scale, we conjecture scaling forms for the correlation and response functions:

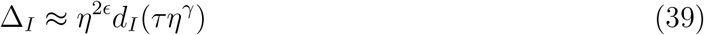

and

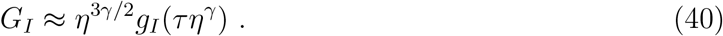

The Lagrange multiplier, *μ*, which parametrizes the properties of the regions of phenotype space that the trajectories are traversing, has dependence on *η* that scales similarly: 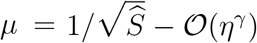 as we expect that the correction to *μ* — which has units of rate — will be of order the inverse of the persistence time scale on which *G*(*τ*) deviates from its marginal form.

To match onto the fast dynamics, Eq.(29), we need that for *x* small, *d_I_*(*x*) ~ *x*^2^. This *τ* dependence implies that the fitness flux *R* ~ *η*^2(*ϵ*+*γ*)^. Similarly, for *x* small, but still *τ* ≫ 1, *g_I_*(*x*) ~ *x*^−3/2^ with the exponent in the prefactor in Eq.(40) forced by the condition that there is no substantial *η* dependence in the fast regime.

To match onto the slow dynamics, for which *G* crosses over to being proportional to ∂Δ, we need *d_I_* ~ *x^θ^* and *g_I_* (*x*) ~ *x*^*θ*–1^ for *x* ≫ 1, but still *τ* ≪ *T_W_*. Together with the 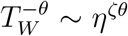 coefficient of the shorter-time end of the slow dynamics, these yield two identities between ζ and the other exponents.

The *η* dependence appears directly in the equation (30) for *R*. If the integral is dominated by large times, then *R* ~ *η*/*T_W_* which, together with the equalities forced by matching to fast and slow times, yields:

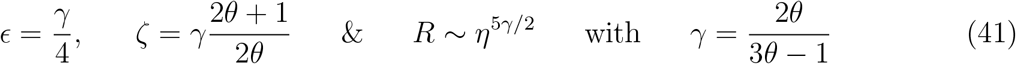

For the pure-*p* model, *θ* = 1, *γ* =1, *ϵ* =1/4 and ζ = 3/2.

The above results are correct for 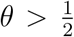 for which the integral for *R* is indeed dominated by large times. But for 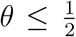, the integral over long times in Eq.(30) diverges as ∫ *dττ*^2*θ*–2^ for *τ* ≪ *T_W_*, so *R* must be controlled by intermediate times: this fixes *γ* = 2 and *R* ~ *η*^5^, but ζ = 2 + 1/*θ* still increases with deceasing *θ* diverging at the transition to the convergent phase.

Thus far, we have *assumed* existence of a solution, on the time scale of the persistence time, of the form Eqs,(39,40), and shown what the conditions for the scaling functions must be to match onto both the fast and slow solutions. It is straightforward to derive coupled quadratic integro-differential equations for *d_I_* and *g_I_* by taking some time derivatives of the full DMFT equations and keeping only the leading terms for small *η*. In principle, one could solve these equations numerically, with *θ* setting the large *x* “boundary” condition and the scale set by the integral condition for *R* which directly involves *η*. We have not attempted to do this as we mainly care that the fast and slow regimes do indeed connect, as claimed, to yield a steady state solution of the DMFT equations even for very small *η*.

#### 4.2.6 Numerics for *p* – *H* model

To check that the Red Queen phase exists for small *η* and indeed has the correlations and response obtained from asymptotic analysis of the presumed steady state, we directly integrate the full mean-field dynamics. We focus on the *p* – *H* model with *p* = 3, for which the critical point is at 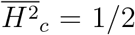 and, for 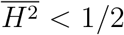, the predicted *C*_∞_(*η* → 0) satisfies 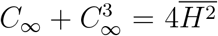 with *β* = 1/(1 + *C*_∞_).

From various initial conditions at *t* = 0, it is straightforward to integrate Eqs.(19,20) with the self-consistency conditions for ϒ and Σ, Eqs.(21,22), as a function of both times, 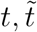, imposed at each time step. Unfortunately, the integro-differential form and the dependence on both *t* and 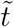 rather than just their difference, means that the integrals have to be carried out at each step which makes it hard to go to the very long times needed to probe the behavior for very small *η*. However for modestly small *η*, 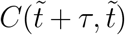 and 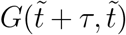 are seen to converge towards functions of just *τ* with the memory of the initial conditions decaying away.

For the pure-*p* = 3 model 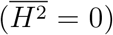 the exponential decay at long times and the predicted relationship between ∂*C*(*τ*) and *G*(*τ*) is readily seen for a substantial range of *η*, with the exponential decay rate, 1/*T_W_*, consistent with the predicted *η*^3/2^ behavior. The fitness flux, *R*, is consistent with the predicted *η*^5/2^ and the intermediate time scale — the persistence time, *τ_persist_* — is indicated by the time at which the maximum of –∂*C* occurs: this is consistent with the *η*^−1^ prediction. The scaling in the intermediate regime as a function of *η^γ^τ* = *ητ* is observed, at least roughly. These results confirm that for the pure-*p* model the short and long time dynamics indeed splice together over an intermediate time range as predicted.

As 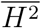 increases, the time scales needed to reach steady state grow — as predicted by the increase of the exponents *γ* and ζ — and it becomes hard to make decent estimates of the exponents from the numerics. Furthermore, the convergence of *C*_∞_(*η*) to its predicted limit as *η* → 0 — eg with *C*_∞_ estimated from *C*(*t*, *t*/2) — is rather slow.

For 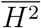 substantially larger than 1/2, exponential convergence to the trivial state with *C*(*τ*) = 1 is seen readily. But the behavior for 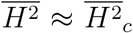 is hard to analyze numerically due to the time scales becoming very long for small *η*, confounded by crossovers from being near a critical point. However a check can be done by looking at the aging behavior for *η* = 0. There is a natural Ansatz for the aging at the critical point, 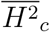: the effects of the slow time scale should be very small as *C*_∞_ ↗ 1, so we conjecture that the intermediate persistence-time scale determines the behavior. This yields, for *η* = 0, *R*(*t*) ~ *t*^−(*α*+2*γ*)/*γ*^ = *t*^−5/2^ at 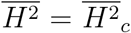. The numerics are quite consistent with this over a wide range of times and deviations from the power law in opposite directions are readily seen for larger and smaller 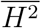, as expected. This is consistent with the prediction 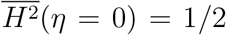 but, as that prediction could be obtained from the breakdown of the single-maximum phase, the correct identification of 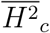 is a less stringent test than are the dynamics and *η* dependencies.

Overall, the numerics support the basic Ansatz’s and predicted scaling behaviors for the dynamics, with the possible exception of the behavior on the slow-time-scales when 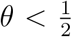 — predicted for 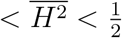 — for which it is hard to get to long enough times. It is possible that in this regime more complicated behavior with extra time scales emerges, as in aging of some physical models. [19] But as our interests are primarily in the existence of the Red Queen phase with small feedback and its qualitative and semi-quantitative properties, we have not pushed the numerics to optimally test the quantitative — but in any case non-universal — predictions.

## 5 Red Queen dynamics from constraints

In the presence of constraints, the DMFT equations for the response and correlation functions have exactly the same form as without constraints but the key self-consistently determined functions 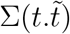 and 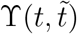 are more complex. The constraint covariance function, 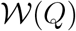, and its derivatives, 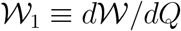 and 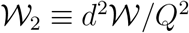, all play a role: note that for 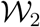 to be non-zero, 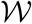 has to be non-linear — quadratic in the minimal model of Eqs.(12,13) — corresponding to the constraint hyper-surfaces being curved.

In addition to *G*, a related response function associated with the constraints plays a role:

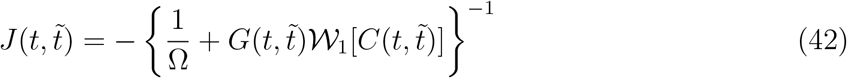

where the inverse is in the sense of an integral equation (equivalent to matrix inverse for discrete time): *J*, like *G*, is causal thus zero for 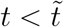. One finds:

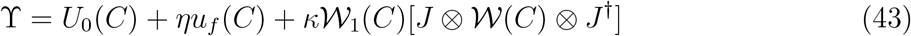

where 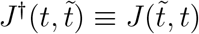, and

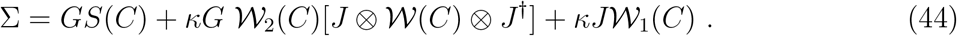

Note that the overall normalization of 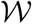 drops out for Ω → ∞ as it must for rigid constraints

### 5.1 Constraints without feedback

#### 5.1.1 Constraints only

In the absence of any environmentally determined contributions to the fitness — *U*_0_ = 0, *S* = 0, *η* = 0 — and if all the constraints can be simultaneously satisfied, there will be a rapid initial transient on time scale of 1/Ω after which a seemingly-trivial steady state obtains. In terms of the DMFT, the stationary state has *C*(*τ*) = 1, Φ = 0, Lagrange multiplier *μ* = 0, and Laplace transform of *G* determined by the coupled equations (20,42,44) with *μ* = 0. As, with *C* constant, the integrals over intermediate times are simple convolutions and the inverse is likewise simple, for the Laplace transforms, 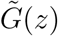 and 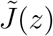, the equations are algebraic. In the rigid constraint limit the linear-response to a sharp kick corresponds simply to unconstrained displacement in a fraction 1 – *κ* of the directions without subsequent relaxation: thus on average we have simply *G*(*τ*) = 1 – *κ* for all *τ* > 0 corresponding to 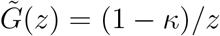.

With soft constraints, but all of them exactly satisfied, at any point on the neutral manifold the Hessian of Φ has *D* – *K* zero eigenvalues and a Marchenko-Pasteur spectrum [23] of *K* negative eigenvalues of order Ω: this is because the rank-*K* stability matrix has the form *γ*^†^*γ* in terms of the gradient matrix: *γ_αj_* = *∂*Γ_*α*_/∂*x_j_*.

But this analysis *assumes* that all the constraints can be satisfied simultaneously. If the constraints are substantially skewed, this can break down. The simplest situation is with just linear constraints 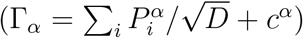 which each restrict **X** to be in a hyperplane that is centered away from the origin for *w_C_* = Var(c*^α^*) > 0. Already for two constraint planes cutting a three-dimensional ball, one sees that if the *c^α^* are too large, the planes need not intersect on the surface. In high-dimensions, random linear constraints become unsatisfiable (with very high probability) above some critical *K* which can be less than *D*: we derive now the general condition for non-linear constraints.

Approaching the unsatisfiabity transition from smaller *K* is difficult because then all times matter so that the correlation and response functions do not depend only on time differences. Thus, instead, we approach by decreasing *K* with soft constraints and check whether the constraints force Φ to remain negative in steady state — and hence are not satisfiable. Assuming *C*(*τ*) = 1 for all *τ*, one finds integrated response

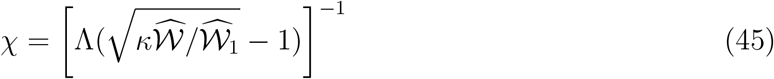

which breaks down as *κ* decreases to a value 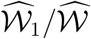 at which point χ diverges consistent with the expectation of 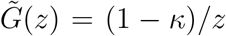 when all the constraints can be satisfied. Thus all constraints can be satisfied only if 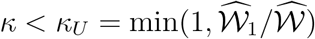.

#### 5.1.2 Three phases with constraints-only

With many soft constraints, there can be a convergent phase with only one maximum or a phase with exponentially many maxima. The condition for stability of the single-maximum DMFT solution is readily found as in Sec. 3.1 (also below): it breaks down for 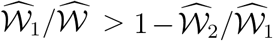. Thus with soft constraints only, there are three possible phases: under-constrained for 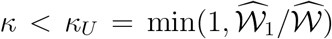, over-constrained but with convergence to a unique negative-fitness maximum for 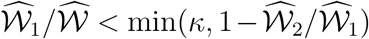, and over-constrained with multiple maxima and aging dynamics otherwise.

#### 5.1.3 Constraints with simple organismic fitness

We now analyze the interplay of constraints and the simplest possible organismic-level environment-dependent fitness: adding to the evolutionary forces a bias, **H**, towards one phenotype. With soft constraints and 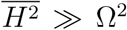, the strong bias from the environment should overwhelm the constraint forces and there will be a unique maximum and simple behavior. As in Sec.3.1 without feedback, we can find the limit of stability of this phase by noting when 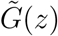 becomes singular at *z* = 0 indicating marginal stability. The static response, χ, and 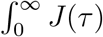 (which are simply related by Eq.(42) when *C* =1 for all *τ*) together determine the stability condition and likewise the condition for the divergent response to an added low-frequency noise. Again, for *η* = 0 both criteria give the critical 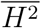. For positive *η*, the latter condition could be used to derive the critical 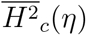 which increases with *η*..

In the rigid constraint limit, there is only one time scale set by 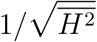, and the system has either one stable fixed point or exponentially many depending only on whether *κ* is greater than or less than a critical value:

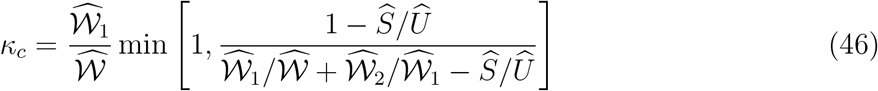

with, again, hats denoting values at *Q* = 1. This expression for *κ_c_* is valid also with more complex fitness correlations as long as 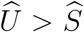. (If 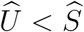 then, as we showed earlier, there are many maxima and Red Queen dynamics with or without the constraints.) For the simplest fitness model with just a tilt to the landscape in the direction of **H**, *S* = 0, so the critical *κ* is determined entirely by the correlations of the constraints with 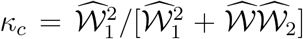. For *κ* < *κ_c_* any tilt in the landscape will be enough for the phenotype to find its optimum — even more easily with soft constraints — while for *κ* > *κ_c_*, sufficiently rigid constraints with stiffness Ω > Ω*_c_*(*κ*) will cause trajectories to wander around, gradually slowing down but never committing to one of the exponentially many maxima.

In addition to the phase boundary at *κ_c_*, for rigid constraints there is also the unsatisfiability boundary (found above) at 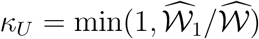. These coincide — for any 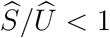 — when 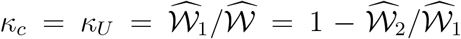. In the parameter space of *κ* and the skewness of the constraints in Eq.(12), which change only 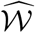, the two phase boundaries intersect at this multi-critical point.

### 5.2 Constraints with feedback

The simplest possible feedback, Sec. 2.3, modifies the organismic level evolutionary forces to

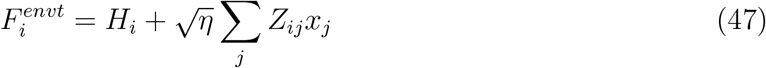

with the elements *Z_ij_* iid gaussian so that 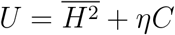 and *S* = 0.

With small feedback, we expect that, as for the simple random fitnome models, from the phase with exponentially many maxima, any amount of feedback will give rise to Red Queen dynamics. The trajectories will wander over all the close-to-neutral manifold on a long time scale, *T_W_*, that is proportional to an inverse power of *η*; on this and longer time scales the fitness flux will approach a non-zero constant.

To analyze the Red Queen phase, we again assume a steady state. First, by taking a derivative of the equation for *C*(*τ*), and using the relationship between *J* and *G*, we obtain exactly the same result for the fitness flux, *R*, Eq.(30), as in the absence of constraints. To find the behavior for small *η*, one can proceed similarly to the case without constraints decomposing both *G* and *J* into fast and slow parts with, on the wandering time scale, the fast parts of both acting like approximate delta-functions in time. The marginality condition in the limit of small *η* can be used to fix the amplitude of these delta-functions, with a negative coefficient for *J_F_*. But in contrast to the case without constraints, some of the long time information is also needed, in particular 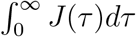, as well as *C*_∞_. On the slow time-scales, one again makes the Ansatz that *G*(*τ*) ≈ –*βdC*_+_/*dτ* and finds that this gives a consistent solution for the pair of equations (19.20). The key identity is that on slow time scales,

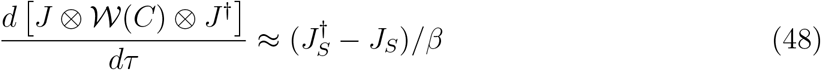

from which follows the needed result that enables relating the *C* and the slow *G* equations,

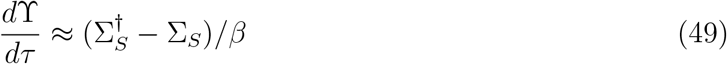

neglecting the terms of order *η*.

By analyzing the large and small *τ*/*T_W_* limits of the slow equations and using the relationship between *J* and *G* as well as the marginality condition for the fast dynamics, a complicated set of relations between the primary undetermined quantities can be derived and used to find, in particular, *C*_∞_, *β*, and the integrated response, χ. In the steady state with memory of the past having decayed away on time scales much longer than *T_W_*, χ (as in Sec. 3.1) determines the average projection of **X** on the direction of **H**. For the particular case of the simplest model with rigid constraints and 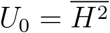, in the limit *η* → 0, χ also determines the average fitness, 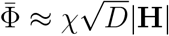.

From this analysis we see that the slow dynamics of the Red Queen phase with constraints and simple feedback are similar to the toy landscape models. The fast dynamics, as probed by the response function and the persistence of trajectories, Sec.4.2.1 are also similar. However the matching between the fast and slow time scales on the intermediate time scale, *τ_persist_*, is somewhat different because of the role of *J*, and we have not analyzed it in detail.

#### 5.2.1 Numerics with constraints and simple feedback

To check that the analytical results with constraints do correctly yield the two phases and boundaries between them as well as the predicted slow dynamics, we integrate the DMFT equations numerically for a specific example. We take the simple feedback model of Eq.(47) so that 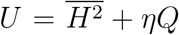 and *S* = 0; soft constraints with Ω = 2 which sets the inverse time scale; constraint correlations 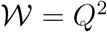; and *κ* = 0.9, considerably above the predicted *κ_c_* = 2/3 from Eq.(46) (which obtains when 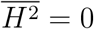 as the softness of the constraints then does not matter). The predicted 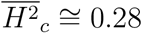 is close to the value at which the long-time aging dynamics without feedback appears most critical. For substantially smaller 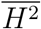, the dynamics and dependencies on *η* are semi-quantitatively similar to those found for the *p* – *H* model with 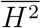 substantially less than 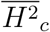, although the range of slow times and values of *η* for which the proportionality of *G* and –∂*C* is observed, is not wide. For 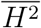 substantially above 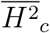, the fitness flux is consistent with going exponentially to zero at long times. While extraction of exponents and confirmation of detailed predictions was not possible from these computations, the consistency of the overall structure, and scalings with time and *η* support the essential similarities between the behaviors of the random fitnome and constraint models.

## 6 Discussion

We have found that in simple high-dimensional models of adaptive dynamics there are two phases. If the complexities of the fitness landscape are overwhelmed by the effects of a strongly optimal phenotype, then the conventional picture of exponential convergence to the fitness maximum obtains. And if modest-strength environmental feedback is added the behavior does not change substantially. With a few distinct maxima, the behavior will be similar except for the initial conditions determining to which fitness optimum an evolutionary trajectory converges.

But if the complexities cause many maxima (and even more saddles) in a high-dimensional landscape, the behavior is very different. In the absence of environmental feedback the trajectories will not commit to any maximum and continue wandering over much of the phenotype space gradually slowing down as an inverse power of time. In such complex landscapes, any small — even simple — feedback from environmental change caused by the evolution will cause Red Queen dynamics with the evolution continuing forever without (after an initial transient) slowing down: in particular, the fitness flux, *DR*, the apparent rate of increase of the fitness relative to the resident population, will approach a non-zero value whose magnitude scales as a power of the strength, *η*, of the feedback. And the deterministically chaotic nature of the dynamics will make the future trajectories highly sensitive to perturbations.

How robust are these predicted behaviors? In particular how general are the existence and properties of the Red Queen phase? We first consider generalizing the class of high-dimensional models and then the effects of various neglected features.

### 6.1 Generality of Red Queen dynamics

The analysis we have carried out already illustrates the generality of the predicted behaviors: the simple random landscapes and the model caricaturing all the complexities by fixed constraints with only simple coupling to the environment, are very different but exhibit very similar behavior. Other landscapes characterized by combinations of high-dimensional gaussian-random functions and low-dimensional structure favoring certain phenotypes — generalizations of the **H** · **X** contribution to the fitness — can be analyzed similarly (and the phenospace need not be a hypersphere). Numerical analysis of the DMFT equations may be needed to determine which phase the system is in and the values of the non-universal exponents that characterize some of the dynamical properties. But the qualitative results should be far more general. And other quantities, such as the Lyapunov exponent of the chaos which parametrizes how nearby trajectories diverge, can also be computed by DMFT.

#### 6.1.1 Doebeli-Ispolatov model

Doebeli and Ispolatov [5] (D&I) have studied adaptive dynamics in a family of models:

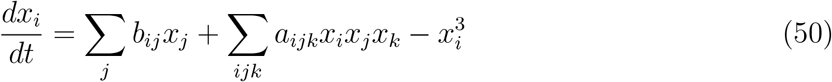

with the {*a_ijk_*} and {*b_ij_*} independently gaussian random with variances chosen (in some of their simulations) to make their effects comparable. If instead of the 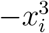 term they had used 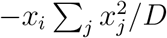, then for large *D* this would effectively confine the trajectories to a hypersphere of diameter set by the variances of random terms, with small fluctuations off the sphere. This would then be in the same class as the models we study except that there are no explicit gradient terms: the corresponding covariances would be *S*(*Q*) = 0 and *U* = *ηu_f_* = *c_b_Q* + *c_a_Q*^2^ with some positive coefficients. In this case, the response function, *G*(*τ*), becomes simply an exponential and a simple expression for the steady-state correlation function, *C*(*τ*), can be derived: it also falls off exponentially for large *τ* and there is only one time scale.

For the model of D&I, DMFT analysis in the limit of large *D* is not straightforward because the equation for *dx_i_*/*dt* has the cubic non-linear term in addition to the gaussian forcing from the interactions with the other components (but no Σ ⊗ *x_i_* term since *S* = 0). A closed form equation for *C* cannot be derived. However the general structure of the equations suggests that again the steady state will be Red Queen dynamics with exponential decay of correlations on a time scale of order unity. The simulations of D&I indeed find that for large *D* the dynamics is chaotic and they estimate the Lyapunov exponent of the chaos and other properties.

#### 6.1.2 Sparse interactions

A key extension is to sparse interactions. The toy model of constraints involved a set of *K* random matrices, 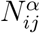, and random vectors, 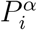, with all elements iid random. However all that is needed for the DMFT analysis is that each component, *x_i_*, is involved in many interactions: with “many” being far less than the *DK* elements of the set of matrix elements of the {*N^α^*} matrices that could affect *x_i_*. Indeed, even if each component is involved in only a few constraints and each interacts with only a few other components, we expect that much of the behavior will be similar. Support for this comes from the study of sparse random matrices with only a few non-zero elements per row and column: the eigenvalue spectrum of these is qualitatively similar but of different shape than the semi-circular form. [24] However such matrices have some number of zero eigenvalues (unless there are more than log *D* elements per row). This would mean, in our context, that some of the components are unconstrained, but this should not matter: those components will quickly go to near whatever value is preferred by the organismic level fitness, then move around a bit driven by the environmental feedback.

Many constraints, each involving only a few of the nano-phenotype components, is a a reasonable caricature of molecular biological constraints — for example on the interaction strengths between protein domains. Some constraints may be better approximated as inequalities — such as Γ_*α*_(**X**) > 0 — than equalities. But combined with the nano-phenotype dependence of the organismic-level fitness, many such inequality constraints together will create a complex landscape with similar properties to the model we have studied. Of course, some inequality constraints may matter in particular environments but not in others: this should not affect the behaviors provided their collective effect does not lessen the constraints so much that the system goes out of the multiple-maxima, or (with feedback) the Red Queen, phase.

One of the advantages of sparse interactions is that it enables simulations to become viable. For example, if in the *p* =3 model only of order *D* (or *D* log *D*) of the *D*^3^ possible elements of the Ξ*_ijk_* set of interactions are non-zero, very high dimensions could be simulated. Similarly, could a model with many soft quadratic constraints of the form Eq.(12).

#### 6.1.3 Nature of environmental coupling and feedback

We have considered the simplest form of the organismic-level fitness and the environmental feedback. More complex organismic level fitness will, as we have shown, enhance the Red Queen tendencies. If the environmental feedback is weak, then the detailed form of it does not matter: all that is required is that the correlations of the forces caused by the feedback are non-trivial, i.e. dependent on **X** (corresponding to *u_f_* (*Q*) depending on *Q*).

The one exception to the independence of the detailed form of the feedback, is if the feedback forces themselves are gradients of some function. An extensively studied example is consumer-resource models [18, 25, 26] in which each strain (or species) consumes multiple difference resources and the consumption of a resource by a strain results in an increase in its growth rate that is — as usually assumed — exactly proportional to the consumption. With the crude caricature of linearizing the effects and responses in **X**, this corresponds to the matrices **A** and **B**^†^ being proportional with a negative coefficient representing the consumption. In this case the matrix **Z** = **AB** is symmetric and its effects are exactly equivalent to an additional term in the fitness 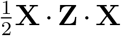 which gives rise to a contribution to *S*(*Q*) that is exactly the derivative of *U*(*Q*). And the same holds for non-linear dependence of the effects and responses on **X**. [26] In these cases, there is no Red Queen dynamics: the total fitness will always be increasing while gradually slowing down. However any small violation of the perfect proportionality between consumption and growth rate — or inclusion of interactions via cross-feeding or non-consumed chemicals (e.g. [27])— will result in a non-gradient part of the forces and again Red Queen dynamics.

### 6.2 Additional features

Several features and processes that we have neglected in the simple model can be understood by analogy with analyses in other contexts. Others are much less well understood.

#### 6.2.1 Stochasticity

Stochasticity of mutations will cause pheno-space evolution to be some form of stochastic gradient ascent. And finite population size will enable occasional steps downhill. Both these effects are very similar to adding thermal fluctuations in the form of Langevin (white) noise added to the equations of motion. (Indeed, in a discrete genotype space, the stochasticity is exactly equivalent to thermal noise in the usually-clonal limit that we consider. [28]) However for dynamics in high-dimensional random landscapes with many maxima, small thermal fluctuations (low temperature) do not change the basic structure of the aging dynamics: like at zero temperature, the average fitness will continue to increase but still slow down as a inverse power of time. Only if the thermal fluctuations are sufficiently large (above a critical temperature) will the behavior change. [16] If small feedback is added at low temperatures, it will still induce the Red Queen phase. [29] Of course the chaos will no longer be deterministic but, given the sensitivity that already exists from the deterministic chaos, the distinction will only be observable on short time scales, roughly, time scales shorter than the persistence time when the deterministic chaos is not yet felt. Extension of our DMFT analysis to include stochasticity is straightforward. [29]

An important point needs making about the difference between the effects of memory-less stochasticity in the dynamics and the effects of the non-gradient deterministic feedback we have studied. One might have expected that once trajectories have “forgotten” their direction after many persistence times, that the feedback terms in the forces would act similarly to memory-less stochasticity. That this is not true is far from obvious! The importance of the memory of past history is reflected in the dependence of properties like the mean fitness flux in steady state, on the whole of the past history, Eq.(30), with a substantial fraction arising from the long time scales, of order *T_W_*, over which the trajectories wander over much of phenotype space. The appearance of an effective “temperature”, 1/*β*, in the relationship between the response and correlations in the Red Queen phase on time scales of order *T_W_*, Eq.(31) does *not* imply that the feedback actually acts like effective thermal noise: the independence of *β* on the feedback strength for small *η* is one indication of this. (See [11] on general issues of effective temperatures.)

#### 6.2.2 Stable islands in the Red Queen phase?

The analysis we have carried out of the Red Queen phase obtains for “generic” initial conditions. In high dimensions, the predicted behavior is expected to occur in all but an exponentially small, in *D*, fraction of the pheno-space: what-one-might-call a “usually Red Queen” phase. However at least for some of the simple models (e.g. *p* – *H* model for small 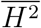) in the absence of feedback there are exponentially many stable maxima with a gap in the eigenvalue spectrum of their Hessian [10] — although collectively their basins of attraction occupy an exponentially small fraction of the space. With small feedback (much less than the size of their gap, – λ*_max_*) most of these maxima should remain stable fixed points. If a trajectory is started sufficiently near to one of these, it will converge exponentially. This has been analyzed for non-gradient models in [29] who indeed found such behavior.

What happens as the feedback strength is increased further? Will the stable fixed points all disappear leading to chaos for all initial conditions? (With the usual trivial exception if started on an unstable fixed points or unstable limit cycle).

Recently Ben Arous et al [12] (henceforth BFK) have shown that for some class of highdimensional random dynamical systems, when the non-gradient parts of the forces are sufficiently strong, all stable fixed points will disappear. They analyzed models of the form *dx_i_*/*dt* = –*μx_i_* + *f_i_*(**X**) with the gaussian forces {*f_i_*} having both gradient of potential and non-gradient parts with covariances sums of *δ_ij_* and *x_i_x_j_* times functions of 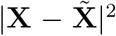: very similar to our forms, and statistically rotationally invariant, but not confined to a hypersphere. The *μ* makes the origin special, and when *μ* (appropriately rescaled) is larger than a critical value, there is a unique stable fixed point. Thus varying *μ* plays a role similar to varying 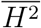 in the *p* – *H* models. And the ratio between the variances of the non-gradient and gradient terms is similar to our *η*. The criteria for instability of the unique stable-fixed-point phase is equivalent to our 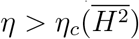 or, equivalently, 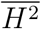 less than a value 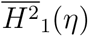 above which there is a unique stable point. For *η* = 0, our 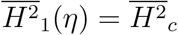, the transition in the absence of feedback. (Note that in BFK’s model, in the stable phase there are no other fixed points, while in ours, because of the topology of the hypersphere, there must be at least one unstable fixed point or saddle in addition to the unique stable fixed point.)

Interpreting BFK’s results in the regime where there are exponentially many fixed points is made more difficult by the quantities they are able to calculate. If *N* is the number of fixed points of a given type — in particular either stable or all — they compute the average 〈*e^N^*〉 over the random {*f_i_*(**X**)}. But this can be dominated by very rare instances in which *N* is anomalously large. However log(〈*e^N^*〉) is an upper bound for the “typical” (or average) log(*N*). Thus if 〈*e^N_stable_^*〉 < 1 this implies that with high probably *N_stable_* is zero. BFK find a regime in which this occurs: in our models, this implies that for 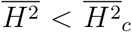, there exists a critical *η*, 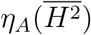, above which there are no stable fixed points and hence always chaotic dynamics: an “always Red Queen” phase. Equivalently, this phase boundary occurs at some 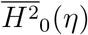. However the boundary that BFK find is *not* equivalent to this boundary, because even if 〈*e^N_stable_^*〉 > 1, it is still possible that *N_stable_* = 0 with high probability. Nevertheless their results translate into an upper bound for 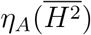. And a lower bound is implied by the stability to small *η* of landscape-maxima that have a gap in their Hessian’s eigenvalue spectrum.

Interestingly, BFK find that the boundary of the convergent unique-stable-point phase — our 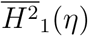 — and the boundary between the usually-Red-Queen and always-Red-Queen phases — our 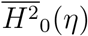 — intersect at *η* = 0. Thus (if their results obtain) for any positive *η*, as 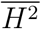 is lowered, we expect the behavior to go *directly* from a convergent phase to an always-Red-Queen phase.

We note that it is possible that for other classes of random landscapes (in particular those with “full replica symmetry breaking”) [10, 19], *any* feedback could destroy all stable fixed points: in such cases, 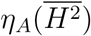 would be zero for all 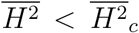, and the usually-Red-Queen phase would not exist. We leave this as an open question.

#### 6.2.3 Finite dimensions

All the results and conjecture we have discussed thus far are in the limit of infinite dimensions: more precisely, they should obtain if all time ranges and parameters such as *η*, etc, do not change with *D* in the limit of large *D*. What are the effects of *D* being large but finite?

In high dimensional random landscapes, the number of maxima is exponentially large in *D*. Generic trajectories will eventually converge to one of the these fitness maxima. As an overwhelming fraction of the maxima are close to marginal, [9, 10] the least negative eigenvalue of their Hessians, λ*_max_*, are expected to have magnitude of order some inverse power of *D*. However it is not clear how the volume of the basins of attraction of the collection of the these dominant maxima (those with λ*_max_* in some small negative range) will scale with *D*: In particular how close to marginal will the ones that collectively attract, say, half the volume, be? Understanding this is needed to predict how the characteristic time and fitness-difference scales depend on *D*. Regardless, the expected behavior of typical trajectories after an initial rapid transient is to slow down as an inverse power of time until a crossover time, *T*_×_, that scales as a positive power of *D*, after which the trajectory will converge to a close-to marginal maximum at a rate of order its very small λ*_max_*. Understanding whether or not –λ*_max_* ~ 1 /*T*_×_, and how these quantities scale with *D*, requires development of better methods: there are few known for high, but finite, dimensional non-linear random systems.

With very small feedback, *η* < *η*_×_ with *η*_×_ scaling as an (unknown) inverse power of *D*, typical trajectories will still converge to stable fixed points at long times. But for *η* ≫ *η*_×_, the basin of attraction of the chaotic Red-Queen strange attractor will occupy most of the volume with the probability of convergence to a maximum becoming exponentially small in *D*.

The model of D&I, Sec. 6.1.1, is qualitatively similar to ours for large *η*. Indeed, they find that in the limit of large dimensions the dynamics are chaotic with very high probability. For the parameters they study in most detail, for *D* = 30 around half of the random parameter values and initial conditions lead to chaos, while by *D* = 80, almost all do. In our models, for small *η*, the dimension above which Red Queen dynamics is extremely likely, is expected to be large — and grow with decreasing *η*. Quantitatively, the dimension needed for chaos with any fixed *η* will certainly depend on the details of the model.

In general, “phase transitions” for infinite *D* will become crossovers for large finite *D*. For example, if a bias **H** · **X** is added with the infinite-dimensional landscape becoming simple above 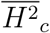, this transition will be smeared out over some range in 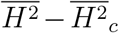 with width of order an inverse power of *D* and, for non-aero *η*, the behavior will be characterized by cross-over functions of 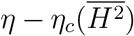 times a power of *D*. For example, the probability that the dynamics is chaotic (over initial conditions and over members of the random ensemble), will be a function of this form.

There do not appear to be published analyses of finite-*D* models in the general class we study except for a recent paper on the matrix model, [21] which derives crossover functions for large *D* and small 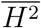 of order an inverse power of *D*.

### 6.3 Evolutionary branching and diversification

The most interesting phenomenon that has emerged from studies of adaptive dynamics is evolutionary branching and diversification.[3, 7] If the evolutionary step sizes are small but not infinitesimal, then it is possible that the resident population and mutant population can stably coexist and the subpopulations each continue to evolve. For models, the only extension that is needed is how the net growth rate of a population of phenotype **Y** depends on the phenotypes and population sizes of the resident phenotypes. A particular simple form of the fitnome, which corresponds to generalized Lotka Volterra interactions, is to assume additive effects of the sub-populations indexed by *a*:

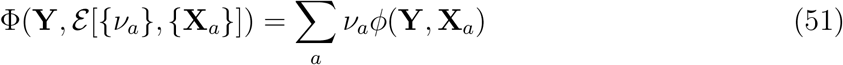

with *ν_α_* the fraction of the population with phenotype **X***_a_* and *ϕ* some complex function. And, of course, one now needs to specify the dependence on **Y** — of, e.g., a mutant — that can be far from all but one of the **X**_*a*_. The condition for coexistence is that for all *a*, 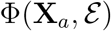 are equal. The matrix of interactions determines both the {*ν_α_*} and the stability, or not, of coexistence on the fast ecological time scale (assumed much faster than the evolutionary time scales). In the spirit of considering the simplest possible models, one could take *ϕ* to be the sum of a part that is independent of the resident populations, and a small feedback part that is bilinear in **X** and **Y** as we have done for the model described in Sec.2.3.

The first question is: Can evolutionary branching occur around the chaotic Red Queen trajectories of the random snowscape models with small feedback? The answer to this is certainly “yes”. But the issue is: How improbable is it for the mutation to go in a direction in which coexistence with its parent can occur? This is addressable by the methods of this paper as it depends on the matrices of the first derivatives of the forces. The next question is more challenging: If the evolutionary trajectories of two coexisting phenotypes are followed will they continue to coexist? And, the most interesting question: Will continual diversification occur leading to a large number of coexisting strains? If this does not occur with the simplest forms of the interactions between the phenotypes and environment, for what forms can it occur? In particular, is a niche-like assumption — i.e. strong decay of the strength of interactions with distance between phenotypes [3, 7] — needed for extensive coexisting diversity to evolve and persist?

While analytic progress on these questions will be challenging, simulations of sparse highdimensional models are viable. Doebeli and Ispolatov have carried out such simulations and find that substantial diversity and evolutionary chaos of the coexisting strains can occur. [7] How general is this likely to be?

A lesson from the analyses of this paper is that key features of high-dimensional models are likely to be independent of many of the details — provided systems are in the same “phase”. Thus, for example, if a “phase” with high coexisting diversity is found in high-dimensional random fitnome models, it is likely to be fairly robust and exist in a wide range of models. Whether or not some niche-like assumptions are needed to induce such a phase, we leave as an important open question.

### 6.4 Connections to reality?

Neither the models we have analyzed, nor the generalizations discussed, are intended to be anywhere near realistic. Nevertheless, one can ask whether the *scenario* of perpetual Red Queen evolution of a mostly-clonal population driven by organismic complexity and weak environmental feedback might be relevant in Nature or the laboratory.

First, should one expect molecular biological constraints on single-celled organisms to lead to over-constrained or under-constrained organisms? On the one hand, if some sets of proteins are over-constrained, duplication and differentiation is a ubiquitous evolutionary process that can get around this and diminish the constraints. On the other hand, if there is pressure on genome size, or generally cost to redundancy — believed the case for bacteria — then cells are unlikely to end up heavily under-constrained. Thus somewhat-under-constrained seems most plausible — and this is the regime in which we find Red Queen dynamics.

Second, is there evidence of — or a good argument for — continual evolution of core proteins driven by selection? Much of protein evolution is said to be effectively neutral with the fixation rate of mutations simply the net neutral mutation rate. [30] However for large microbial populations with typical coalescent times more than 10^7^ generations ago, “effectively neutral” is a very stringent condition. Surely not all observed seemingly-neutral protein evolution is neutral enough! Yet in large microbial populations, any mutation that is unconditionally beneficial would have fixed long ago. Thus non-neutral mutations that fix in a species or sub-population are most likely conditionally beneficial or deleterious — even if only very slightly — depending on genomic background, environment, etc. One key property that all proteins must maintain is to not interfere substantially with others: in particular not binding to them too strongly. Thus whenever one protein changes driven by substantial — and perhaps “interesting” — selective pressures, many others will feel weak selective pressures to change slightly so that they do not bind to the modified protein. Of course, there are likely to be many amino acids that could change in order to do this, and the selective pressure will be time dependent because of ongoing evolution — at least on long time scales. Thus disentangling what drives what, which changes are effectively neutral, etc, is impossible. But this basic argument already suggests that Red Queen evolution driven by complex onstraints and weak feedback via the environment is a highly plausible process for driving continual evolution on both long and short time scales.

We have argued that, even with this *potential* mechanism for continual evolution, there are two “phases” that could occur: the Red Queen phase, and a convergent phase in which trajectories converge to one of a modest number of “fitness maxima” — more properly, stable fixed points — even with environmental feedback. Why might long-term biological evolution lead to populations that undergo Red Queen rather than convergent dynamics? The simplest argument is that populations in the Red Queen phase will maintain genetic diversity and flexibility to change in response to changes in environmental pressures: These are likely to survive better in the long run than populations that converge to a stable state. And the earlier argument about tendency for the number of functional constraints to increase but not too far, also suggests that Red Queen dynamics might be much more ubiquitous — at least in microbial populations — than adaptation that approaches a fitness peak.

Finally, we raise the question of possible relevance for experiments. Recent analyses of the long term *E. coli* evolution experiments of Richard Lenski’s group, indicate a rather surprising behavior at late times.[31] After a rapid initial transient over a few thousand generations. the rate of fixation of mutations and how fast they fix — which depends on their selective advantage — slows down but then seems to not change much over the subsequent 60,000 generations. [31] In contrast, the increase in fitness relative to the ancestor seems to continually slow down. [31] It is clear that some ecological effects play a major role in these evolutions. But is it possible that gradual evolution of the environment driven by the continual evolution of the bacteria induces as well more subtle changes that drive the evolution to continue without appreciable slowing down? Although one could not reasonably argue that these experiments support the scenario developed in this paper, the ubiquity of the Red Queen dynamics that we have found should change one’s view of what is — or more importantly what is *not* — surprising.

## Acknowledgements

The work was supported by the National Science Foundation via PHY-1607606.

